# Dissociable effects of LSD and MDMA on striato-cortical connectivity in healthy subjects

**DOI:** 10.1101/2025.02.07.637042

**Authors:** Natalie Ertl, Imran Ashraf, Lisa Azizi, Leor Roseman, David Erritzoe, David J Nutt, Robin L Carhart-Harris, Matthew B Wall

## Abstract

**Introduction:** Lysergic acid diethylamide (LSD) and 3,4-Methylenedioxymethamphetamine (MDMA) are widely used psychoactive drugs and their potential use in psychiatric medicine is currently generating interest. The mechanism by which these drugs may assist recovery in addiction, mood disorders and post-traumatic stress disorder (PTSD) is still not well understood. Most investigations of the effects of these drugs on brain activity have focussed on cortical resting-state networks, however the striatum is a key reward and motivation hub of the brain and aberrant striatal processing may be part of the pathophysiology of these disorders. Consequently, we investigated striatal connectivity following acute MDMA and LSD administration.

**Method:** Resting-state fMRI (rs-fMRI) data were acquired, and seed-voxel functional connectivity analyses were used with the striatum subdivided into three seed regions: the associative, limbic, and sensorimotor striatum. Within-network connectivity was measured using group mean network maps and whole-brain connectivity (seed-to-voxel) was also examined.

**Results:** Neither MDMA nor LSD significantly changed within-network connectivity of any of the three striatal seed regions. However, striatal connectivity with other brain regions was significantly altered with both MDMA and LSD. Most notably, MDMA reduced connectivity between the limbic striatum and the amygdala, while LSD increased connectivity between the associative striatum and the frontal, sensorimotor, and visual cortices.

**Conclusion:** Changes in connectivity were mostly observed outside the standard striatal networks, consistent with previous findings that psychedelics reduce network modularity or between-network segregation and increase connectivity across standard networks.

## Introduction

In recent years, the intersection between recreational substances and psychiatric medicine has garnered significant attention, particularly in the case of psychedelics such as Lysergic acid diethylamide (LSD), psilocybin, and N,N-Dimethyltryptamine (DMT), as well as the entactogen 3,4-Methylenedioxymethamphetamine (MDMA). The use of psychedelics as medicines has been practised by different groups around the world for thousands of years (Hardy 2021), however their recreational use and the consequent legal restrictions has held back scientific research and clinical applications for decades (Doblin et al. 2019; Dyck 2006; Nutt, King, and Nichols 2013).

Classic psychedelics are agonists at serotonin 5-hydroxytryptamine (5-HT)_2A_ receptors (Nichols 2016), causing profound cognitive disturbances and mood-altering effects (Johnson et al. 2019). LSD is a classic psychedelic with a particularly long duration of up to 12 hours (Hwang and Saadabadi 2023). Its powerful impact on perception and consciousness has made it a subject of extensive research (Carhart-Harris et al. 2016; Roseman et al. 2016; Dyck 2006; Levine and Ludwig 1965) with users commonly reporting enhanced sensory perceptions, vivid hallucinations, and alterations in the sense of self (Schmid and Liechti 2018). LSD was widely used clinically in the first wave of psychedelic research in the 1950s and 1960s (Fuentes et al. 2020) for numerous disorders including alcohol addiction (Dyck 2006; Smart and Storm 1964; Abramson 1966), depression (Savage 1952), and management of pain and anxiety in terminal illness (Schimmel et al. 2020; Pahnke et al. 1970). Its extended pharmacokinetics have made it a less popular choice for modern clinical research, with most studies preferring shorter-acting compounds such as psilocybin, however one recent study has shown promising results for treating anxiety (Holze et al. 2023).

Early neuroimaging studies with LSD showed increased blood flow and connectivity with the visual cortex under LSD, which correlated with subjective reports of complex visual imagery (Carhart-Harris et al. 2016). Other studies have identified increases in global functional connectivity, and these effects have correlated with LSD induced ego dissolution (Tagliazucchi et al. 2016), and other subjective effects (particularly in the somatomotor network; Preller et al. 2018). Functional connectivity changes also interact with music listening and correlate with eyes-closed visual imagery (Kaelen et al. 2016). Graph theory analyses have demonstrated that LSD produces more network functional complexity, increases randomness, and decreases segregation of functional brain networks (Luppi et al. 2021). In common with other classic psychedelics such as psilocybin (Petri et al. 2014) and DMT (Timmermann et al. 2023), LSD therefore has a wide-scale disruptive effect on cortical brain networks, and tends to increase network integration and decrease segregation/modularity.

MDMA (sometimes now called midomafetamine) is an entactogen that acts as a monoamine releaser and a reuptake transporter inhibitor (Lizarraga et al. 2015). This makes it more akin to amphetamines (Gough et al. 1991), with only mild hallucinogenic properties. Like classic psychedelics, its primary impact is on serotonin (5-HT), however MDMA’s stimulant-like qualities comes from its lesser effects on dopamine and noradrenaline (Liechti et al. 2000). Unlike typical stimulants, it functions as an empathogen, enhancing social-emotional processing (Carlyle et al. 2019). MDMA was researched in therapeutic settings in the late 1970s and early 1980s, predominantly for couples therapy and trauma management due to its empathogenic properties (Greer and Tolbert 1986). The neuroimaging research into MDMA’s effects on the brain are limited in comparison to LSD, however task-fMRI studies have suggested MDMA increases frontoparietal activation in the Go/No-Go task (Schmidt et al. 2017), may alter the amygdala responses to emotional faces (Bedi et al. 2009, though see Schmidt et al., 2018 for conflicting results) and has effects on the response to emotional autobiographical memories (Carhart-Harris et al. 2014). Functional connectivity analyses show that MDMA likely does not have the wide-spread disruptive effect on resting-state networks characteristic of classic psychedelics (Roseman et al. 2014) but seed-based analyses have implicated connectivity changes in the medial temporal lobe (amygdala, hippocampus) and insula as being implicated in the action of MDMA (Carhart-Harris et al. 2015; Walpola et al. 2017). Like LSD, MDMA is currently being investigated for its potential therapeutic use, for a number of disorders (Sessa, Higbed, and Nutt 2019).

Most existing neuroimaging research on LSD and MDMA has primarily focused on the effects of the drugs on cortical network systems, or sub-cortical regions such as the thalamus (Preller et al. 2018). However, many of the psychiatric conditions for which these substances are being investigated, such as PTSD and addiction, are thought to be rooted in striatal dysfunctions (Yager et al. 2015). The striatum, a collection of subcortical regions, can be functionally divided (Martinez et al. 2003; Tziortzi et al. 2013a; Joel and Weiner 2000) into three key areas: the limbic striatum, which includes the nucleus accumbens and the inferior portion of the putamen; the sensorimotor striatum, encompassing the superior portion of the putamen and the tip of the caudate; and the associative striatum, which covers the remaining parts of the caudate and putamen. These subdivisions are critical for various functions: the limbic striatum is involved in motivation, reinforcement, and emotion, the sensorimotor striatum in habit formation and motor learning, and the associative striatum in decision-making and cognitive control (Martinez et al. 2003).

The striatum is highly innervated by the substantia-nigra and the ventral tegmental area, receiving over 80% of all the dopamine neurons in the brain. The distinct functional subdivisions are each involved in different cortico-striatal loops, and so play a significant role in many psychiatric disorders that might be amenable to treatment with either LSD or MDMA. For this reason, we investigated changes in striatal connectivity using resting-state fMRI data from acute challenge studies with these two agents.

## Methods

The current data are reanalyses of previously published research on MDMA (Carhart-Harris et al. 2015) and LSD (Carhart-Harris et al. 2016); please see these publications for the full study protocol.

### Study design

#### LSD

The LSD study used a single-blind (participant blinding only), balanced-randomised design. 75µg LSD or placebo was administered via identical IV solutions (more information in table 2). Participants attended two scanning days at least two weeks apart to minimise carry-over effects, previous studies have reported a variable half-life of 75µg LSD, suggesting it ranges from 2.2 to 4.3 hours (Family et al. 2022). Patients lay inside a mock MRI scanner for 60 minutes and were encouraged to relax with their eyes closed, in order to acclimatise the participants to the MRI scanning environment and to reduce any potential anxiety later in the trial. Following this, subjects were moved to the real MRI scanner for a set of scans which included: a structural scan, arterial spin labelling (ASL) and resting-state fMRI. Two resting-state scans were completed per treatment session, one at 70mins post dose and the next at 90 mins, separated by a music listening scan. Participants were encouraged to lie with their eyes closed. After the MRI scanning, there was a break of approximately 35 minutes, after which magnetoencephalography (MEG) scanning was performed (not reported here). Once the subjective effects of LSD had sufficiently subsided, the study psychiatrist assessed the participant’s suitability for discharge.

**Table 1.**
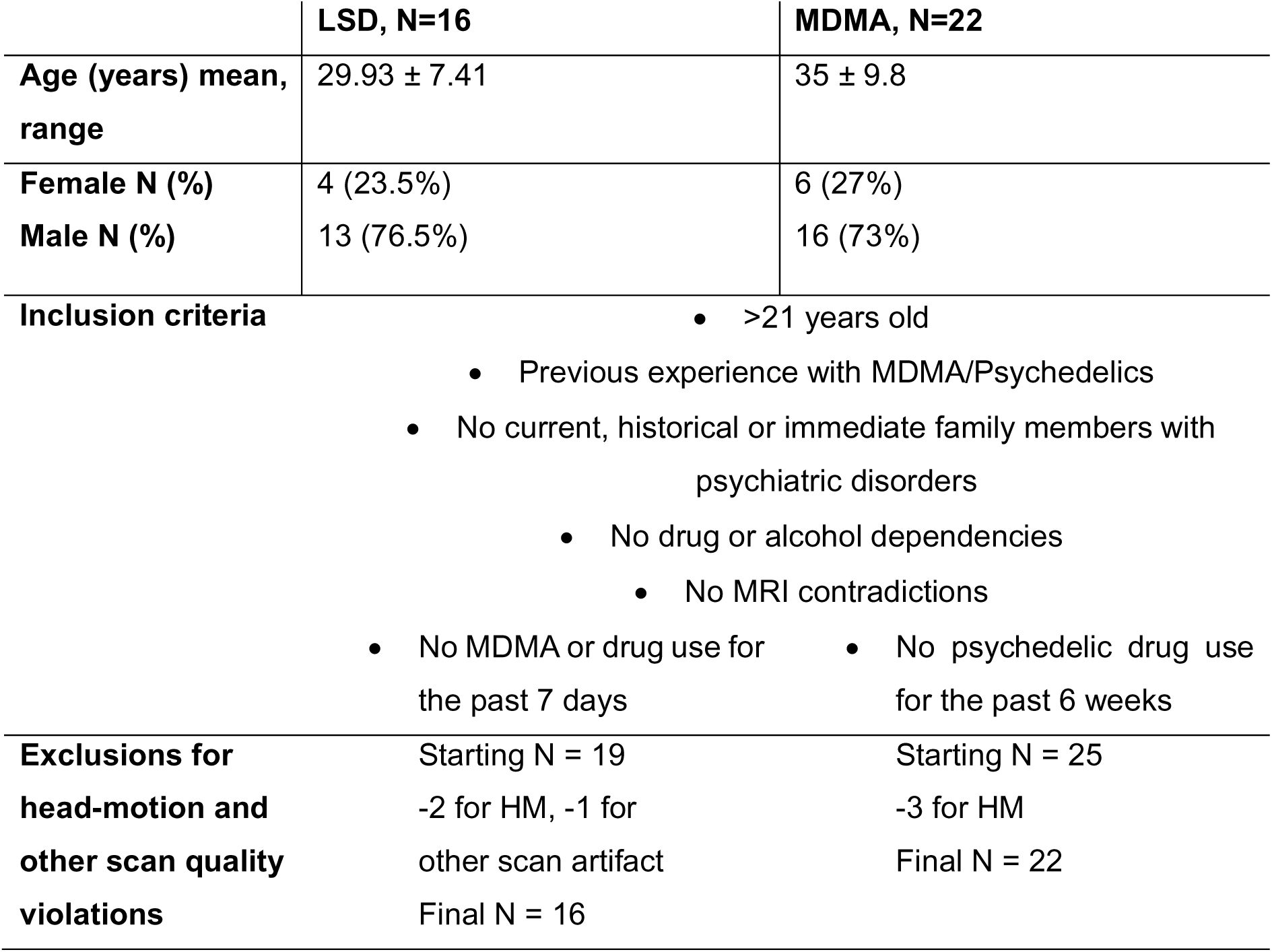
Participant characteristics for LSD and MDMA study and main inclusion criteria.

**Table 2.** Summary of MRI acquisition of the two studies.

| **Functional resting-state scans** | **LSD** | **MDMA** |
| --- | --- | --- |
| **Scanner** | 3T GE HDx system | 3T Siemens Tim Trio scanner |
| **Sequence** | T2*-weighted echo-planar images (EPI) | T2*-weighted echo-planar images (EPI) |
| **Repetition time (TR)** | 2s | 2s |
| **Echo time (TE)** | 35ms | 31 ms |
| **Isotropic voxel size** | 3.4mm | 3mm |
| **Slice acceleration factor** | SENSE = 2 | GRAPPA = 2 |
| **Flip angle (degrees)** | 90 | 80 |
| **Axial slices per TR** | 35 | 36 |
| **Total scan time** | 7 min 20 sec | 6 min |

#### MDMA

The MDMA study was a double-blind, placebo controlled, within-subjects, randomized-controlled trial. The participants underwent two scanning sessions, seven days apart, identical other than the administration of either MDMA or placebo in a randomised order. Both participants and researchers were blinded to the substance administered. The administration route was via identical capsules (see table 2 for more details). Participants underwent two resting-state scans per visit (four in total). The first scan took place approximately 60 minutes after drug administration and the second at approximately 113 minutes post drug administration. Participants were encouraged to relax in the MRI machine with their eyes closed.

#### Participants

#### Drug administration

LSD was administered intravenously. 75µg LSD in 10ml saline was administered over two minutes, followed by saline wash. The placebo was 10ml saline, administered over two minutes, followed by saline wash. MDMA was administered orally. The dose was a 100mg capsule of MDMA-HCL, while the placebo was 100mg of vitamin C.

#### MRI acquisition

Anatomical T1 weighted images were acquired in both scans. The details of the functional acquisitions are listed in table 3.

#### fMRI analysis

All analyses were conducted using FMRIB Software Library (FSL) 6.0, and broadly following an approach used in previous work (Ertl et al. 2023; Wall et al. 2019; Ertl, Freeman, et al. 2024; Wall et al. 2022; Demetriou et al. 2018; Comninos et al. 2018). The data was pre-processed using standard procedures: spatial smoothing with a 6mm FWHM (full-width, half-maximum) Gaussian kernel, high-pass temporal filtering (100s), head-motion correction using MCFLIRT and non-linear registration to a standard template (MNI152). At this stage head motion parameters were examined, and each scan was assessed for mean framewise displacement >1mm or maximum displacement >3mm in any direction, if these measures were exceeded on either session, then the participant was excluded. Framewise displacement measures were derived using the fsl_motion_outliers function and paired *t* tests on these values were conducted between the placebo and drug conditions, in order to check for head-movement differences between the treatments.

The anatomical data were parcellated using FMRIB’s automated segmentation tool (FAST), producing white matter (WM) and cerebrospinal fluid (CSF) segmentations. These were coregistered into individual subjects’ functional data space and thresholded at 0.5. The mean time-series from these parcellations were extracted to be used as nuisance regressors in the model (Power et al. 2014).

Seed-voxel analysis was used to assess striatal functional connectivity; this approach assumes that voxels which are activating in a similar manner to that of the seed are likely functionally connected. The striatal seeds selected followed the original parcellation by Martinez et al. (2003), using the atlas provided by Tziortzi et al. (2013), and are shown in supplementary figure 1. Each seed (in MNI152 space) was registered to the participants structural and then functional scan. The time-series from the participants’ seed regions was then extracted and this was used as the regressor of interest in the model. The WM and CSF regressors, along with an extended set of head-motion parameters (set of 24 head-motion parameters, including six original regressors: three translations, three rotations, plus temporal derivatives and quadratic versions of the original six regressors) were also added to the model.

Next fixed effects mid-level analyses were run to average the results from the first and second scan for each treatment session of each participant. These mid-level analyses were then used in the higher-level models. All group-level analyses were conducted using FMRIB’s local analysis of mixed effects (FLAME-1) method, employing cluster-level thresholding (*Z* = 2.3, *p* < 0.05) to account for multiple comparisons. This threshold provides appropriate control for Type I errors when used with FSL’s FLAME-1 model (Eklund, Nichols, and Knutsson 2016), while also maintaining sensitivity and thereby minimising Type II errors (Slotnick 2017; Noble, Scheinost, and Constable 2019). Initially, as a validation step, a group mean (all subjects, all scans) analysis was performed to produce overall networks for each seed region, these networks were corroborated against previous studies (Ertl et al. 2023; Wall et al. 2019; 2020; Ertl, Freeman, et al. 2024). To investigate overall within-network connectivity, these derived networks were thresholded at 50% of their maximum Z values and binarised to produce network masks (see figure 1). A mean connectivity parameter estimate for each participant for each drug condition was extracted from these masks and a paired *t* test was conducted to investigate significant differences in within-network connectivity between placebo and drug scans.

**Figure 1.**
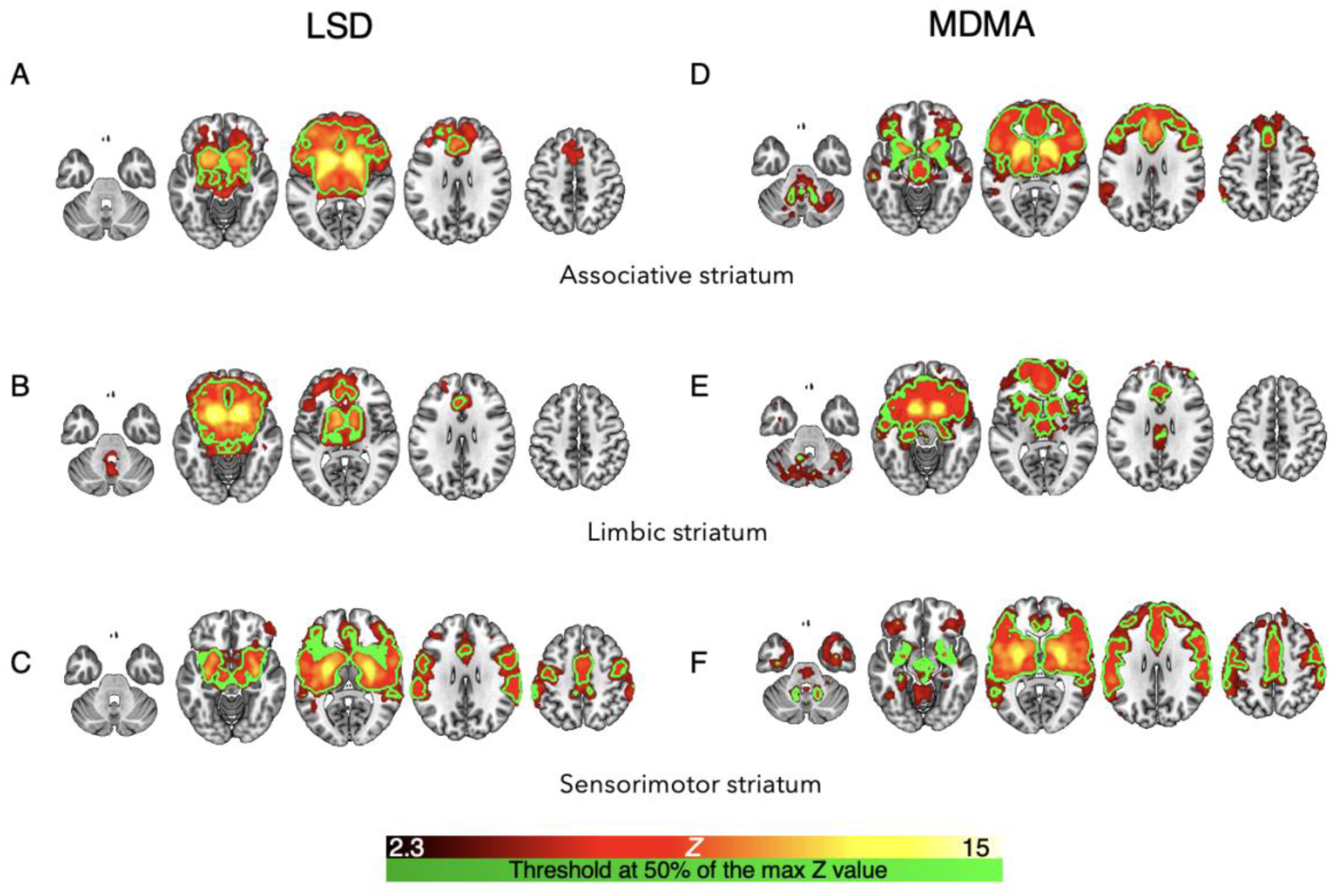
Mean network maps averaging placebo and LSD with the A) associative striatal seed, B) limbic striatal seed, C) sensorimotor striatal seed; and placebo and MDMA with the D) associative striatal seed, E) limbic striatal seed, F) sensorimotor striatal seed. LSD N=16, MDMA N=22, results are cluster corrected (Z=2.3) and thresholded P<0.05. Overlayed in green outline is the region thresholded at 50% of the max Z score mask used to derive the network ROI in the subsequent analyses.

Next, to test for changes in connectivity with each network and the rest of the brain, seed-voxel analysis was performed with a within-subjects effects model. This showed regions of the brain which were relatively more or less connected with the seed regions in the active drug compared to placebo scans.

## Results

### Head-motion

Two participants were excluded for excessive head-motion (>3mm max frame-wise displacement) and one for a separate artifact, from the LSD analysis; once they were removed from further analysis, a paired *t* test showed a significant difference in mean framewise displacement between the placebo and LSD condition *t*[31]=4.58, P<0.0001). Three participants were excluded from the MDMA analysis due to excessive head-motion, there was also a significant difference in mean head movement between the placebo and MDMA conditions *t*[43]=2.29, P=0.026.

### Within-network connectivity

Striatal connectivity networks were derived from averaging across the placebo and drug conditions for each seed region. These networks closely match previous work using different subject cohorts (Ertl et al. 2023; Wall et al. 2020), with the associative striatum characteristically showing connectivity with the frontal lobe, the limbic striatum being strongly connected with medial-temporal-lobe regions, and the sensorimotor striatum showing strong connectivity with the motor cortex. Similar patterns of activation were seen in the LSD (figure 1A-C) and MDMA groups (figure 1D-F), thereby validating this seed-voxel approach and the analysis procedures. The network ROI (derived from thresholding at 50% of the maximum Z value), is also shown on figure 1 as a green outline. Group average results of the placebo and drug conditions are separately shown in supplementary figure 2.

Changes in connectivity within the derived network ROIs were then investigated. The networks were thresholded to 50% of the maximum Z value and binarized to produce network masks (green outline in figure 1). Connectivity parameter estimates were then extracted from these masks to provide connectivity measures from within the network in placebo and active drug administration. No significant changes in within-network connectivity were found in any of the striatal seeds with LSD or MDMA administration (Figure 2)

**Figure 2.**
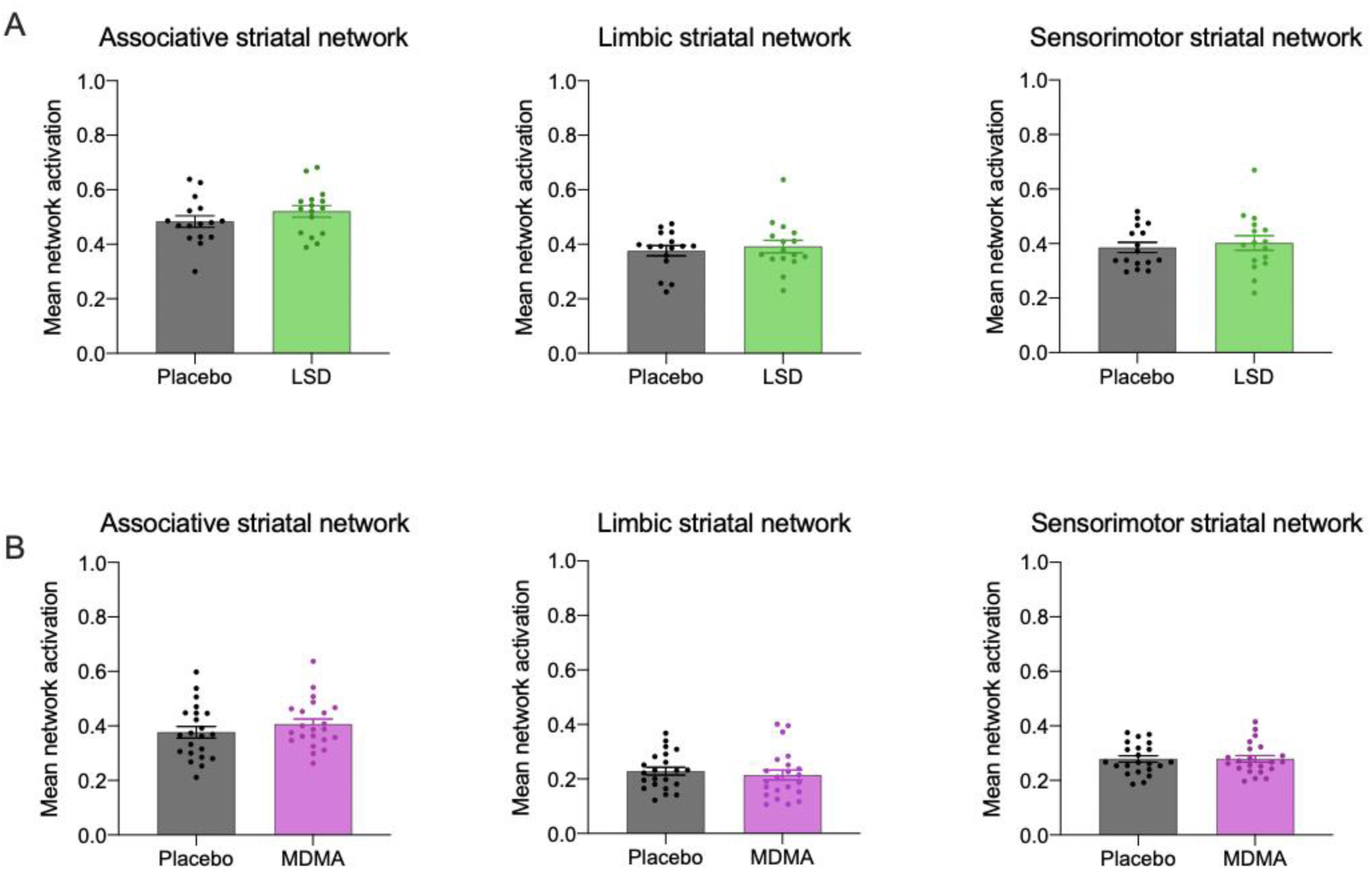
No changes in within-network connectivity were identified in any of the striatal networks with A) LSD (green) or B) MDMA (pink) administration relative to placebo, LSD N=16, MDMA N=22, error bars show SEM.

### Seed-voxel (whole-brain) analysis

Next, striatal network connectivity with the rest of the brain (whole-brain, seed-to-voxel analyses) was investigated.

#### LSD

Acute LSD administration altered whole-brain connectivity with all three of the striatal seeds investigated (Figure 4A-C). Increases in connectivity were observed between the associative striatum and a large cluster spanning the visual cortex, as well as the bilateral sensorimotor cortex and medial frontal regions. Decreased connectivity was observed with the tail of the left putamen (part of the sensorimotor striatum) extending into the thalamus (Figure 3A). The limbic striatum significantly increased its connectivity with the left frontal orbital cortex and an area around the inferior occipital cortex from the occipital pole extending into the lingual gyrus. Decreased connectivity was observed with an area in the extending from the inferior parietal cortex into the superior lateral occipital cortex (Figure 3B). Finally, connectivity with the sensorimotor striatum and an area in the bilateral parahippocampus extending into the temporal cortices and an area around the precuneus extended into the intracalcarine cortex significantly increased under acute LSD. A decrease was observed around the rostral anterior cingulate and the left inferior frontal gyrus (Figure 3C).

**Figure 3.**
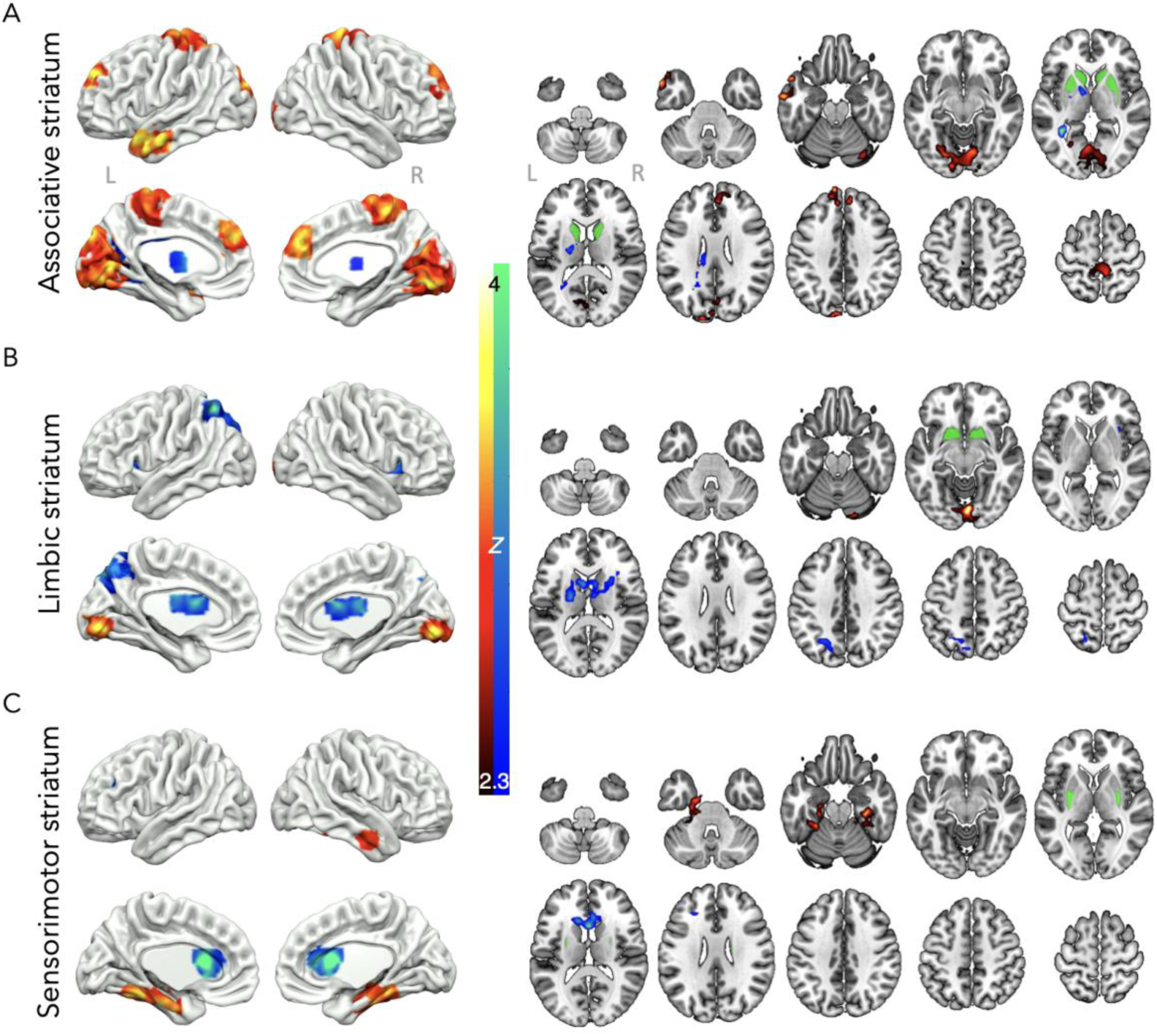
Acute LSD significantly increased (red/yellow) and decreased (blue/green) connectivity between the A) associative striatum, B) limbic striatum, and C) sensorimotor striatum and areas in the rest of the brain, N=16, results are cluster corrected and thresholded at *Z*=2.3, *p*<0.05, original seed regions shown in green on axial slices.

**Figure 4.**
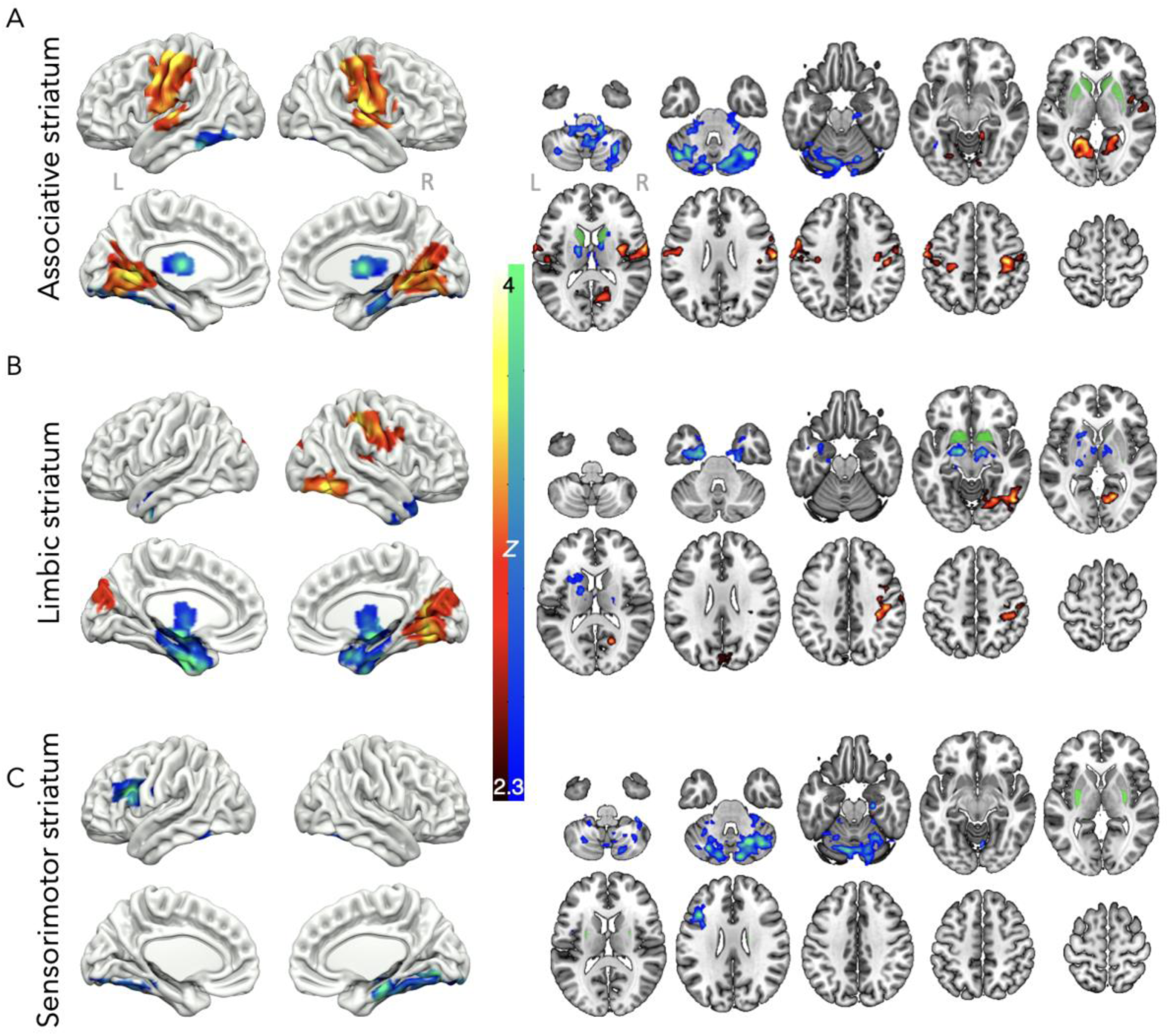
Acute MDMA significantly increased (red/yellow) and decreased (blue/green) connectivity between the A) associative striatum, B) limbic striatum, and C) sensorimotor striatum and areas in the rest of the brain. Results are cluster corrected and thresholded at *Z*=2.3, *p*<0.05, N=22, original seed regions shown in green on axial slices.

#### MDMA

MDMA also significantly altered brain connectivity with all three of the striatal networks investigated (Figure 4, A-C). Under acute MDMA treatment, associative striatum connectivity with the sensorimotor cortex and lingual gyrus was significantly increased, whereas connectivity with the cerebellum was significantly reduced (Figure 4A). Connectivity between the limbic striatum and the sensorimotor cortex as well as higher order visual areas such as the lingual and intracalcarine cortex was also increased, while connectivity between the limbic striatum and the thalamus, amygdala, and hippocampus was significantly reduced under acute MDMA administration (Figure 4B). Finally, connectivity between the sensorimotor striatum and the inferior frontal gyrus and cerebellum was reduced under acute MDMA administration (Figure 4C).

## Discussion

The results show that neither LSD nor MDMA have strong effects on the broad measure of within-network connectivity relative to placebo. However, both drugs produced marked changes in connectivity with brain regions outside of the standard striatal networks. This aligns with previous research suggesting that psychedelics may reduce brain modularity (R. Daws et al. 2021) and increase global functional connectivity (Preller et al. 2020; 2018). When modularity is reduced, overall network structure is degraded, and brain regions/networks become more interconnected and communicate more with other regions/networks that they typically do not synchronise with. This effect of psychedelics may allow for novel patterns of thought and perception to emerge, and be related to their clinical effects (Daws et al. 2022). Previous work has shown strong effects of the classic psychedelic psilocybin on cortical between-network connectivity, with relatively few effects of MDMA (Roseman et al. 2014). In contrast, here we show that MDMA may actually have reasonably comparable effects to the classic psychedelic LSD on striatal networks, though with important distinctions as well.

### LSD

Of the three striatal networks investigated under acute LSD administration, the associative striatum showed greatest changes in connectivity, with significant increases in the lingual gyrus, sensorimotor cortex, frontal poles, and the orbital cortices, as well as a small reduction in connectivity with the thalamus. The associative striatum has functions in cognitive integration and memory processes (Grill et al. 2024). The lingual gyrus and medial occipital lobe are involved in visual processing (Palejwala et al. 2021) and increased connectivity with these areas under LSD may reflect the intense visual hallucinations often reported with an LSD experience. Aqil and Roseman (2023) have recently emphasised that the unique effects of psychedelics on visual function are an important (and perhaps, overlooked) part of their phenomenology and that low-level sensory effects are likely to significantly influence higher-level brain systems. The finding here that the (‘cognitive’) associative striatum’s connectivity with the visual cortex is strongly increased with LSD supports this thesis. Under acute LSD the associative striatum increased connectivity with the sensorimotor cortex, possibly enabling deeper and more meaningful sensory-emotional experiences and a heightened sense of feeling connected to surroundings or others (Forstmann et al. 2020). Additionally, increased functional connectivity was observed between the associative striatum and the frontal poles. The frontal cortex is involved in executive control and cognition (Friedman and Robbins 2022), so this change in connectivity may reflect the alterations in cognitive function often experienced under acute LSD, for example increased reinforcement learning ability (Kanen et al. 2022).

Relative to the associative striatum, LSD caused less disruption in limbic striatal connectivity. The limbic striatum contains the nucleus accumbens which has a prominent role in reward and consequently addiction; the limited changes in connectivity observed with these addiction-relevant areas is consonant with the lack of addictive properties seen with classic psychedelics such as LSD (Das et al. 2016; Nutt, King, and Phillips 2010). Moderate changes in limbic striatal connectivity were observed with the visual cortex again emphasising the altered connectivity of the visual cortex during an LSD trip likely leading to hallucinations or altered visual perception.

The sensorimotor striatum is involved in the integration of sensory information and motor planning and execution (Bizzi and Ajemian 2020). It receives inputs from various sensory modalities, processes this information, and is involved in the coordination and execution of voluntary movements (Tewari, Jog, and Jog 2016). Additionally, the sensorimotor striatum is essential for habit formation, procedural learning, and motor skill acquisition (Gremel and Lovinger 2017). Under LSD administration, increased connectivity was observed between the sensorimotor striatum and regions in the temporal lobes including the parahippocampus. This may be significant since the parahippocampus is involved in spatial memory (Bohbot et al. 2015). LSD also reduced connectivity between the sensorimotor striatum and the left inferior frontal gyrus, this reduction in connectivity with the brain’s inhibitory control centre under acute psychedelics may be reflective of the feelings of euphoria experienced under psychedelics as well as the sense of dissociation or altered bodily awareness. Users of psychedelics often report feelings of mental lightness, disembodiment, or a disconnection from physical boundaries (Roseman et al. 2014).

### MDMA

Under acute MDMA administration, increased connectivity between both the associative and limbic striatum with the sensorimotor cortex was observed. The associative striatum is pivotal in cognitive processing (Grill et al. 2024), while the limbic striatum also has a role in emotional processing as well as reward/motivational systems (Pavuluri et al. 2017), and so this increased connectivity with the sensory integration brain regions could reflect the enhanced emotional experiences under MDMA (Hysek et al. 2014). Interestingly, reduced connectivity was observed between both the associative and sensorimotor striatum with an area in the cerebellum under acute MDMA. The cerebellum is also involved in motor coordination and reduced connectivity with this area may reflect the changes in balance and motor coordination seen with MDMA (Stephenson et al. 1999). Specifically, the cluster of deactivation spanned the cerebellar vermis bilaterally. Since the vermis is a key area in oculomotor control (Manto et al. 2012), decreased connectivity with this area could represent the reduced eye-motor control and increase in saccadic movements sometimes experienced with MDMA users (Dumont et al. 2007).

Limbic striatal connectivity with the amygdala and parahippocampus was downregulated under acute MDMA administration. MDMA’s inhibitory effect on the amygdala is particularly noteworthy, given the amygdala’s central role in fear response and fear memory formation (Frick et al. 2022). Individuals with PTSD often have hyperactivity in the amygdala (Morey et al. 2012) and have exaggerated fear responses, hindering their ability to engage effectively in therapy (van der Kolk 2000). By attenuating the amygdala’s response, MDMA could enable individuals to access their trauma memories in a controlled manner, facilitating therapeutic interventions without triggering excessive fight or flight reactions. These findings are consistent with previous work showing changes in amygdala connectivity are related to symptom improvement in PTSD patients treated with MDMA therapy (Singleton et al. 2022). Moreover, evidence suggests that survivors of the 7^th^ October attacks in Israel who had taken MDMA showed some protection against the development of PTSD compared to survivors who had not taken MDMA (Netzer et al. 2024), suggesting MDMA may actually prevent the traumatic memories from forming in the same way, potentially through a relative functional disconnection involving the amygdala.

Acute MDMA administration led to a decrease in connectivity between the sensorimotor striatum and a region around the left inferior frontal gyrus and a region in the cerebellum. Specifically, this cerebellar deactivation cluster spanned the right vermis and crossed into the right Crus I and a small area in the left vermis, involved in motor and oculomotor control (Manto et al. 2012). The left inferior frontal gyrus is thought to be part of the brain’s inhibitory control network (Liakakis, Nickel, and Seitz 2011). Connectivity between the inferior frontal gyrus and sensorimotor striatum may play a role in suppressing unwanted or inappropriate motor responses. A reduction in connectivity between these regions might imply a temporary disruption in the brain’s ability to regulate and inhibit certain movements.

### Clinical Implications

Altered striato-cortical connectivity is observed in addiction and affects cognition, working memory, attention, and decision-making (Gould 2010). LSD may enhance connectivity between the associative striatum and frontal cortex, which could improve cognition and self-control, and so reduce impulsivity. This may suggest potential in treatment for addiction (Zafar et al. 2023), though further research is needed. MDMA, known for its effects on emotional openness and enhancing social interactions, may reduce inhibitory control, allowing emotional release. Its increased connectivity between the limbic striatum and sensory cortex may help process trauma, supporting its potential in PTSD treatment (Mitchell et al. 2023). However, scientific rigor in MDMA research needs improvement (Reardon 2024).

### Strengths and limitations

This study is among the first to specifically investigate changes in striatal connectivity under classic and atypical psychedelics. The placebo-controlled design accounts for between subject differences, allowing for a more accurate detection of the effects of the drugs on striatal connectivity to be uncovered. Despite rigorous correction for head-motion and exclusion criteria, significant differences in head-motion between drug and placebo sessions remain, meaning that head-motion cannot be ruled out as a potential confounder. Mitigating against this interpretation though are the null findings on within-network connectivity, where head-motion might also be expected to have an effect (if it was a potentially serious confounder). While these are re-analyses of some of the largest LSD and MDMA research studies to date, they are both still relatively small sample sizes in fMRI research. Hopefully as psychedelic research becomes more mainstream and legal restrictions are conceivably relaxed, future studies should aim to replicate the present analysis with larger sample sizes. Moreover, the distribution of males to females in both studies is unbalanced. Research is currently suggesting that psychedelics may have particular benefits to women, which have yet to be studied in a clinical setting (Stein et al. 2022; Argento et al. 2022). Future studies should aim to make recruitment for trials such as these more appealing to female volunteers.

## Conclusion

In conclusion, our study examined the neurobiological effects of one classic, and one atypical (LSD and MDMA) psychedelic, revealing nuanced alterations in striatal connectivity and broader brain networks. Notably, MDMA exhibited a reduction in connectivity between the limbic striatum and amygdala, suggesting a functional mechanism for the therapeutic intervention in PTSD. Meanwhile, LSD increased connectivity between the associative striatum and frontal and visual cortices, shedding light on the drug’s impact on sensory integration, creativity, and altered cognition. The impact of this data suggests LSD may have potential as a therapy in reward-based disorders where the brain states need to be remapped, such as addictions (Zafar et al. 2023), obsessive compulsive disorder (Buot et al. 2023), or hypoactive sexual desire disorder (Ertl, Mills, et al. 2024). These findings may underscore the mechanism by which psychedelic medicines enhance emotional intelligence, empathy, and creative expression. By unravelling the neural intricacies of MDMA and LSD on striatal connectivity, our research adds to the body of literature which will hopefully enable targeted therapeutic applications, promising hope for individuals grappling with addiction, trauma, and emotional challenges.

## Supplementary material

**Supplementary Figure 1.**
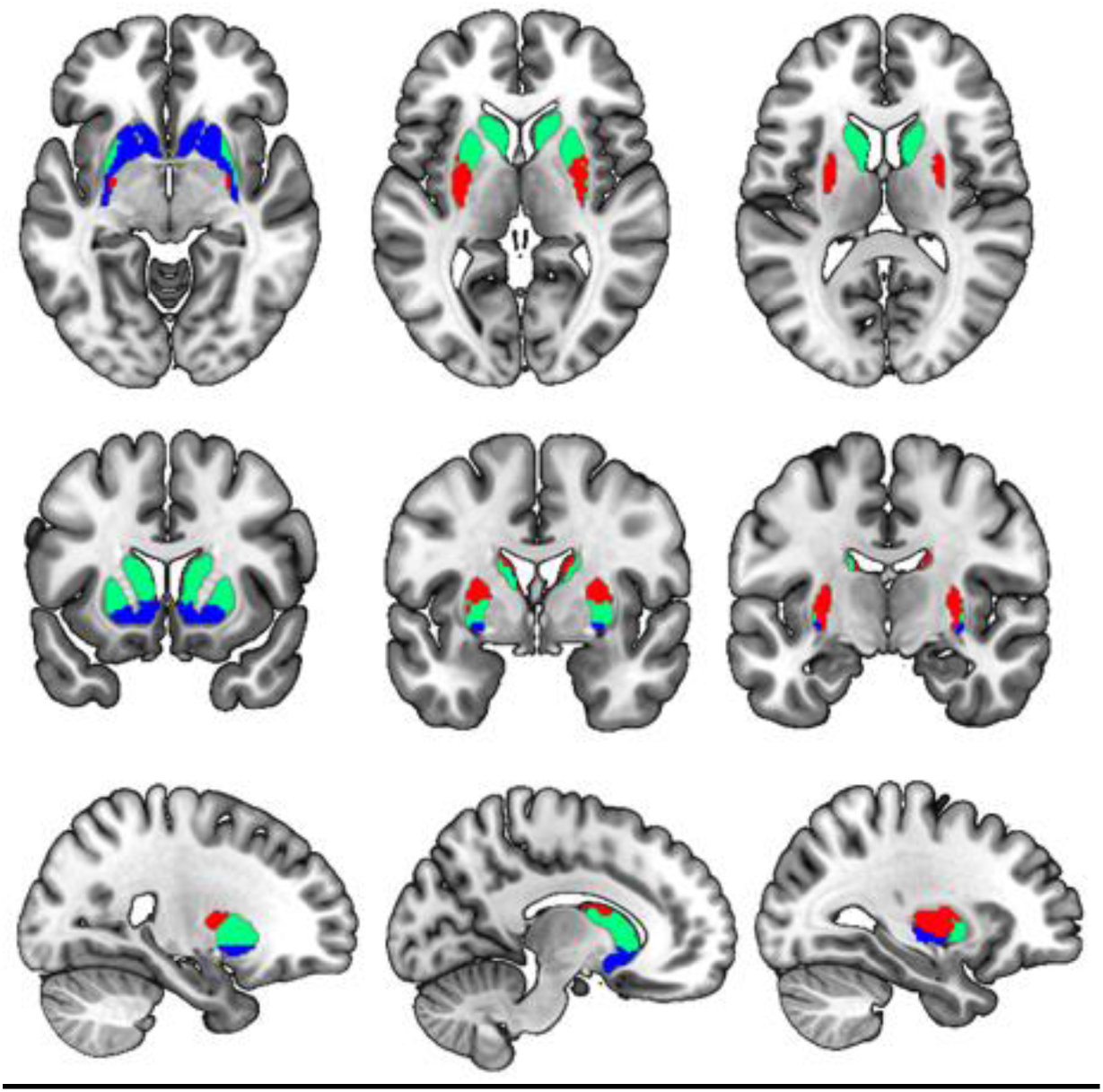
The striatal networks were defined using the associative (green), limbic (blue) and sensorimotor striatum (red).

**Supplementary Figure 2.**
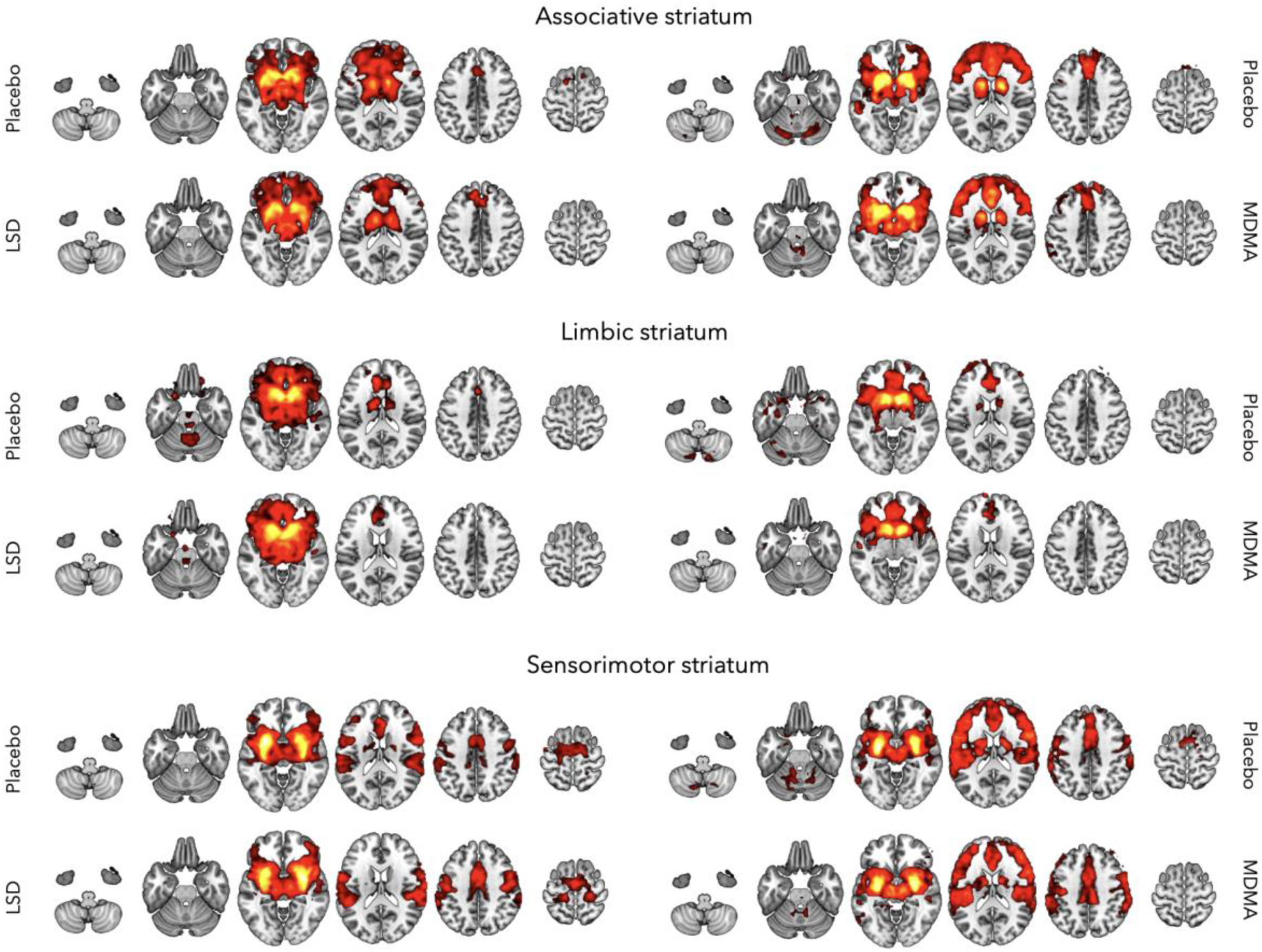
Group average of placebo and drug conditions separately.

